# Molecular Dynamics Study of α-Synuclein Domain Deletion Mutant Monomers

**DOI:** 10.1101/2024.03.23.586267

**Authors:** Noriyo Onishi, Nicodemo Mazzaferro, Špela Kunstelj, Daisy A. Alvarado, Anna M. Muller, Frank X. Vázquez

**Affiliations:** Department of Chemistry, St. John’s University, Queens, NY 11439, USA

**Keywords:** intrinsically disordered proteins, α-synuclein, molecular dynamics, monomer, inter-domain contacts

## Abstract

Aggregates of misfolded α-synuclein proteins (asyn) are key markers of Parkinson’s disease. Asyn proteins have three domains: an N-terminal domain, a hydrophobic NAC core implicated in aggregation, and a proline-rich C-terminal domain. Proteins with truncated C-terminal domains are known to be prone to aggregation and suggest that understanding domain-domain interactions in asyn monomers could help elucidate the role of the flanking domains in modulating protein structure. To this end, we used Gaussian accelerated molecular dynamics (GAMD) to simulate wild-type (WT), N-terminal truncated (ΔN), C-terminal truncated (ΔC), and isolated NAC domain asyn protein variants (isoNAC). Using clustering and contact analysis, we found that removal of the N-terminal domain led to increased contacts between NAC and C-terminal domains and the formation of interdomain β-sheets. Removal of either flanking domain also resulted in increased compactness of every domain. We also found that the contacts between flanking domains in the WT protein result in an electrostatic potential (ESP) that may lead to favorable interactions with anionic lipid membranes. Removal of the C-terminal domain disrupts the ESP in a way that could result in over-stabilized protein-membrane interactions. These results suggests that cooperation between the flanking domains may modulate the protein’s structure in a way that helps maintain elongation and creates an ESP that may aid favorable interactions with the membrane.

## Introduction

Large aggregates of α-synuclein (asyn), known as Lewy bodies, are one of the key indicators of Parkinson’s disease (PD) and form by the oligomerization and fibrilization of misfolded asyn proteins[1-6]. Although asyn is strongly associated with PD, it plays a crucial role in the regulation of synaptic vesicle trafficking and neurotransmitter release in neuronal cells[7-10]. Experimental evidence also suggests that asyn helps control synaptic vesicle fusion by reducing the vesicle curvature[11]. Investigations of the binding of asyn to phospholipid micelles have found that the binding leads to distortion and flattening of the micelle surface[12]. When interacting with a vesicle, the N-terminal region of asyn adopts an amphipathic α-helical structure in membranes that can lead to membrane remodeling[13-15], while the rest of the protein extends to the cytosol.

Asyn is an intrinsically disordered protein (IDP) that can adopt many conformational changes in solution and in cellular environments[16-18]. The wild type (WT) protein consists of three domains: an amphipathic N-terminal domain, the hydrophobic non-amyloid component (NAC) domain, and the acidic and proline-rich C-terminal domain. The NAC domain is the part of the protein most associated with toxic aggregation and fibril formation[16,17,19-21]. Domain truncated asyn can occur in cellular environments due to proteases, lysates, or even splice variants[22]. Some of these domain deletion mutations are known to accelerate aggregation and fibrilization[21-33]. Deletion of the C-terminal domain, specifically, is strongly associated with aggregation and increased fibrilization[22-31]. C-terminal deletion has also been shown to form polymorphic fibrils with twisted Δ-sheets[24,27]. Within Lewy bodies, most asyn proteins show a truncated C-terminal domain, and removal of the domain has been shown to increase the propensity for aggregation[23,34]. Additionally, recent work has found that the human appendix, even in healthy patients, contains C-terminal truncated asyn monomers and various forms of oligomers[25], likely due to the lysates found in the organ.

The N-terminal domain controls the interactions between asyn and the membrane[32], and the KTKEGV imperfect repeat units in the domain have been shown to inhibit the formation of Δ-sheet structures and the formation of fibrils[21]. Studies of fibril and nucleation kinetics have found that residues 1-36 on the domain are fibril inhibiting, while residues 37-60 enhance fibril formation[28]. Removal of the N-terminal domain was also shown to form a polymorphic asymmetric Δ-rich fibril[33].

The flanking domains seem to serve a protective role against monomeric toxic aggregation, but the specifics of how they regulate protein structure are not fully understood. Previous studies have demonstrated that the N-terminal and C-terminal regions of α-synuclein engage in transient intramolecular interactions, which play a crucial role in modulating the protein’s conformational ensemble and aggregation propensity[16,19,20,35-38]. To study how interactions between flanking domains are affected by domain deletions, we have used all-atom Gaussian accelerated molecular dynamics (GAMD) simulations[39,40] to understand how the removal of the flanking domains affects inter-domain contacts, secondary structure, compactness, and electrostatic potential. From our structural analysis and studies of contacts, we found that the N-terminal domain can form contacts between both the C-terminal domain and NAC domain simultaneously. We also found that removal of the N or C -terminal domains leads to increased compactness in the NAC domain, which can indicate increased toxicity. Contacts between the N and C -terminal domains in the WT proteins can lead to an electrostatic potential (ESP) that may help the protein regulate its interactions with anionic lipid membranes and other proteins. Removal of the N-terminal domain disrupts the positive electrostatic potential and removal of the C-terminal domain leads to increased positive charge that could over stabilize protein-membrane interactions. From these results we propose that both domains may work cooperatively through inter-domain contacts to maintain relative elongation in all three protein domains and modulate the electrostatics of the protein.

## Results and Discussion

### N-Terminal Domain Forms Contacts with NAC and C-Terminal Domains

The full sequence of the wild type (WT) asyn protein is shown in Figure 1a with the amphipathic N-terminal domain (residues 1-60) colored green, the hydrophobic NAC domain (residues 61-95) underlined and colored orange, and the acidic, proline rich C-terminal domain (residues 96-140) colored blue. To understand the role of the flanking domains, we simulated the WT protein and three domain deletion variants (Figure 1b): an N-terminal domain deletion (ΔN), a C-terminal domain deletion (ΔC), and an isolated NAC domain (isoNAC).

**Figure 1.**
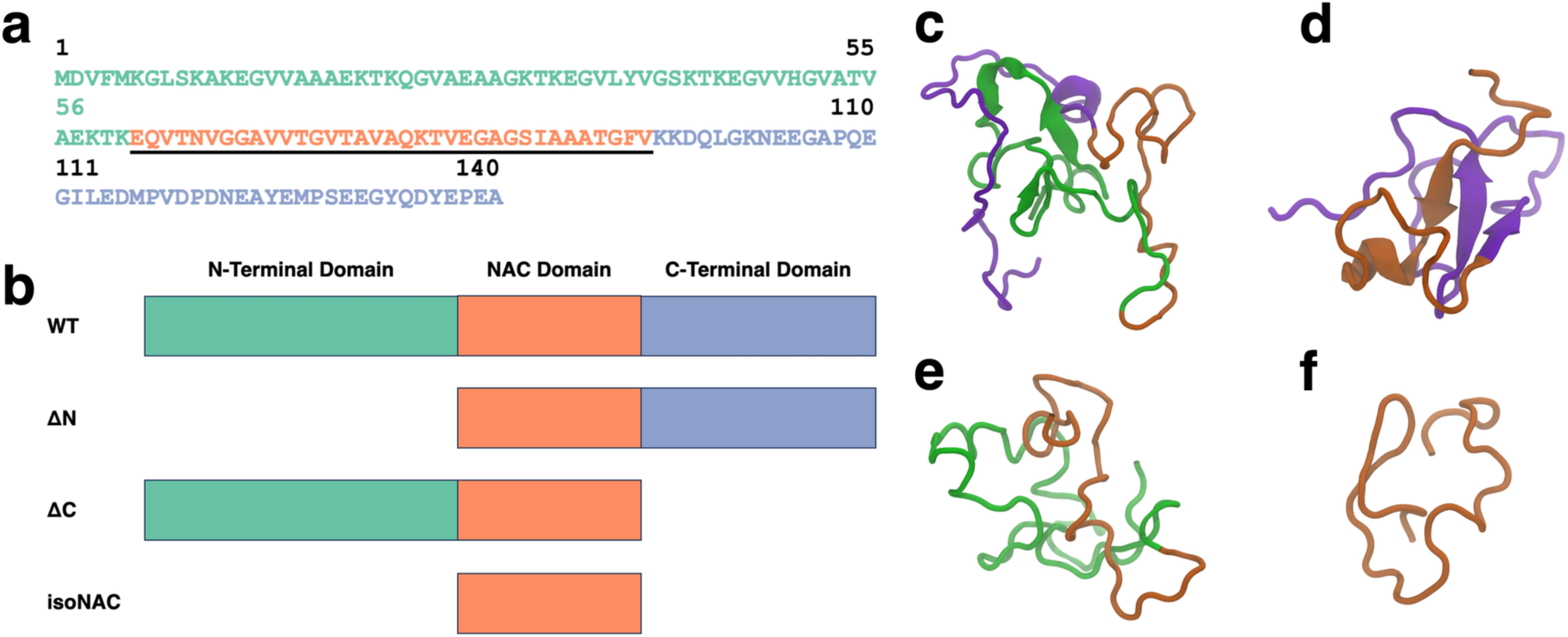
The full WT-asyn protein has three domains: N-terminal, NAC, and C-terminal domains. The protein sequence (a) shown in this figure is colored to highlight each domain, with the N-terminal domain shown in green, the center NAC domain underlined and orange, and the C-terminal domain colored blue. Four variants were simulated for this study (b): WT, ΔN, ΔC, and isoNAC. The representative structures for the most occupied clusters of the WT (c), ΔN (d), ΔC (e), and isoNAC (f) are shown using the New Cartoon representation with the N-terminal domain in green, the NAC domain in orange, and the C-terminal domain in violet.

We used all-atom molecular dynamics (MD) to run our simulations. Standard, unbiased molecular dynamics simulations are generally unable to sample many conformational changes, and this limits their ability to study an IDP such as asyn. Disordered protein MD simulations usually require some sort of enhanced sampling method to explore the folding landscape. We used GAMD to run our simulations[39,40]. The benefit of this method is that it does not require predefined collective variables for biasing and can correctly reproduce the shape of the potential landscape, while still allowing the protein to sample the entire folding landscape. It should be noted that GAMD does not preserve the relative well depths without reweighting but does preserve distribution minima and maxima and their relative rankings[39-42]. The primary goal of our study was to compare the relative differences in structural properties and conformational sampling between the domain deletion variants and the wild-type (WT) protein. Since we are not interpreting absolute free energy values, but instead focusing on qualitative comparisons of the sampled states, the reweighting procedure was not required. This approach was well-suited for our analysis, as GAMD’s ability to identify relative conformational preferences and dominant structural ensembles remains robust even without reweighting[41,42].

We used k-means RMSD clustering[43] to understand the conformational states adopted by each of the asyn variants. The representative structures of the largest cluster for each of the WT (Figure 1c), ΔN (Figure 1d), ΔC (Figure 1e), and isoNAC (Figure 1f) variants shows qualitatively the types of interactions that can occur between the flanking domains. In the WT structure, we found that the N-terminal domain places itself between the C-terminal and the NAC domains, forming contacts with both the C-terminal and NAC domains. In the ΔN structure, we see significant contacts formed between the C-terminal and NAC domains. Additionally, the structure shows the formation of a large β-sheet motif between the two domains. This is in contrast with the WT structure, which has some β-content but only in the N-terminal domain. The representative structure for the ΔC structure still shows contacts between the N-terminal and the NAC domains but does not show the same β-sheet motifs seen in the WT and N-terminal domains. Lastly, the representative structure for the NAC domain shows a conformation that appears to be relatively compact.

Similar structures where the N-terminal domain forms contacts with both the C-terminal and NAC domains are also seen in the other cluster center structures. The representative structures from the top ten WT (Figure 2a), ΔN (Figure 2b), and ΔC (Figure 2c) clusters were aligned to the NAC domain and are shown together (top) along with the surface representation of the most occupied cluster center (bottom). The overlayed structure for the WT protein shows that the N-terminal domain consistently forms contacts with both the C-terminal and NAC domain. This is further exemplified in the surface representation, where the contacts can be directly observed in the structure. Once the N-terminal domain is removed, the C-terminal and NAC domains appear to increase the contacts between the two domains. The surface representation also shows there are more contacts being formed between the two domains than in the WT protein. Removal of the C-terminal domain is more curious. It appears that removal of the C-terminal domain affects the specificity of the contacts. In the WT and ΔN variants, the contacts with the NAC domain seem to show specificity with respect to the region where they form. In the ΔC variant, the N-terminal domain does not seem to be as localized with respect to the NAC domain as in the other two variants. The surface representation shows there are contacts between the N-terminal and NAC domains, but part of the surface area of the N-terminal domain is exposed. Additionally, we aligned the representative structures of the isoNAC variant (Figure S1). We found that all the isoNAC conformations form relatively compact structures, though some were more elongated than others.

**Figure 2.**
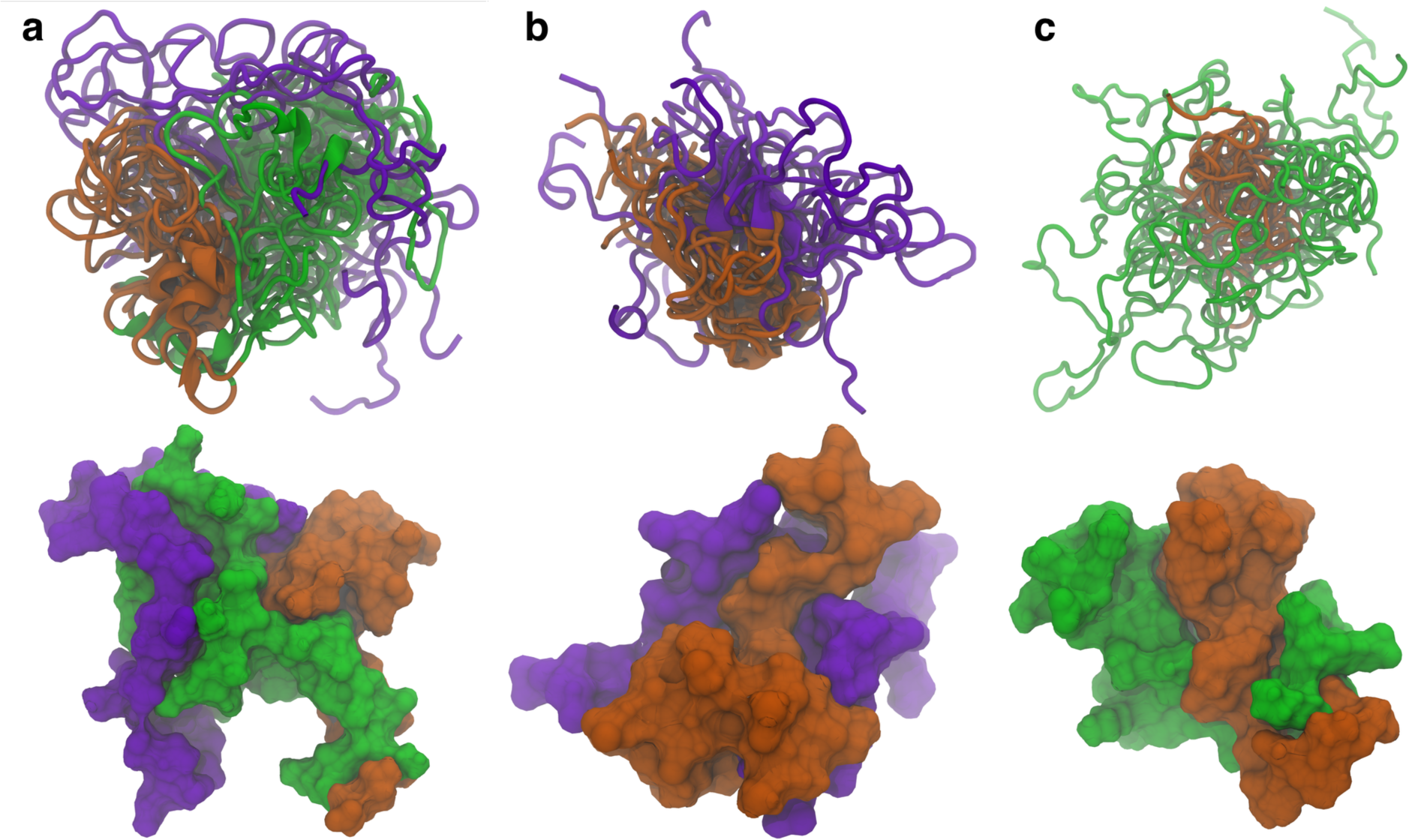
The representative structures of the top ten clusters for the WT (a), ΔN (b), and ΔC (c) variants are shown together (top) along with the SASA representation of the representative structure of the most populated cluster (bottom). The structures are all aligned with respect to the trace of the NAC domain. The overlaid structures are shown using the New Cartoon representation. The N-terminal domain is shown in green, the NAC domain in orange, and the C-terminal domain in violet.

To understand the types of contacts formed between the domains in the WT protein, we calculated the distribution of overall contacts (Figure 3a), hydrophobic contacts (Figure 3b), hydrogen bonds (Figure 3c), and salt bridges (Figure 3d) for the protein. A contact between residues was measured if the distance between C_α_ atoms was less than 6 Å. The overall contacts show that the WT protein sampled structures with more NAC and N-terminal domain (NAC-N) contacts or N-terminal and C-terminal (N-C) contacts than NAC and C-terminal domain (NAC-C) contacts, though there were still structures sampled with a significant number of NAC-C contacts. In general, though, most of the sampled WT structures had few NAC-C contacts compared to NAC-N or N-C contacts. When we further investigated the types of contacts that were formed, we found that the NAC-N interactions involved more hydrophobic contacts than the NAC-C or N-C interactions. On the other hand, the N-C interactions were dominated by hydrogen bonds and salt bridges. The NAC-N interaction, though, also seemed to involve some hydrogen bond formation. The N-C contacts appeared to be the only interactions with a significant amount of salt bridge formation. This result appears to be consistent with experimental studies that have found that long-range electrostatic interactions form between the N-terminal and C-terminal domains in WT asyn[16,19,20,36]. Additionally, disruption of the electrostatic contacts between the N and C domain has been implicated in the formation of asyn fibrils and is associated with unfolding of the monomer during the fibril seeding process[38].

**Figure 3.**
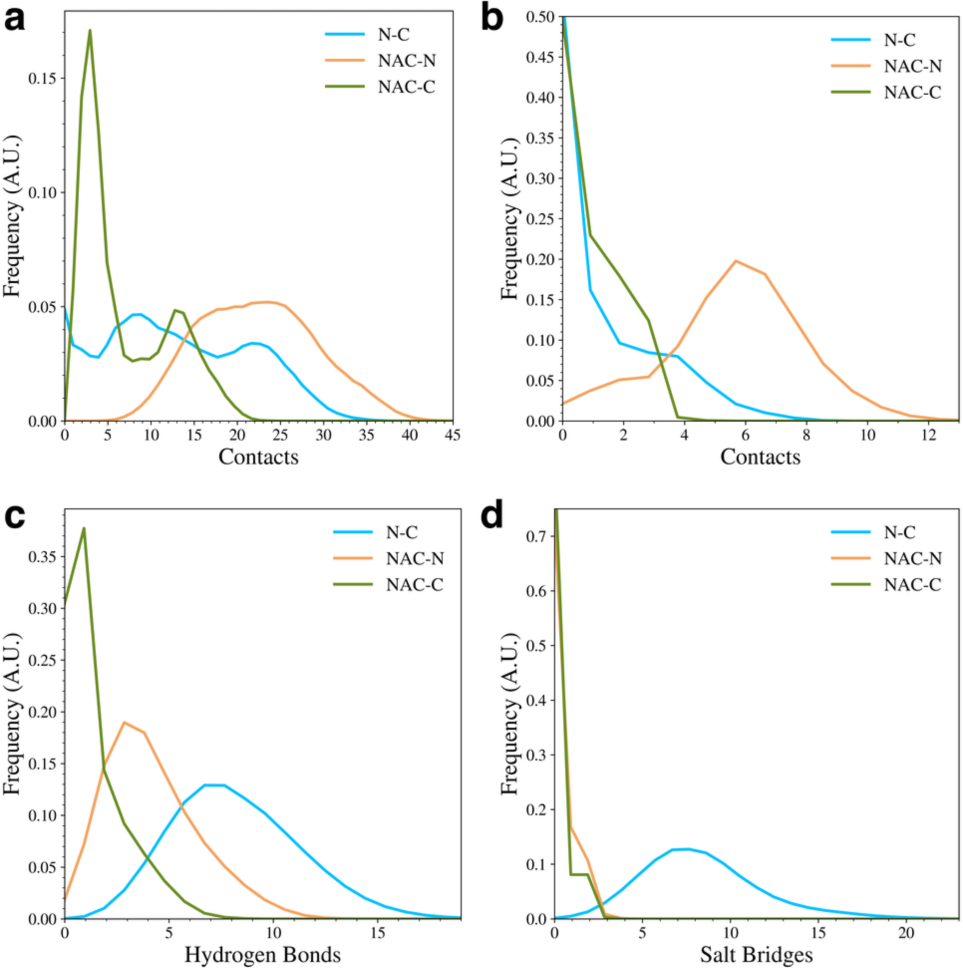
The distributions of interdomain contacts (a), hydrophobic contacts (b), hydrogen bonds (c), and salt bridges (d) in the WT protein.

The contact analysis further highlights that the N-terminal domain can form contacts with both the NAC and C-terminal domains simultaneously because of its amphipathic character. The non-polar region of the N-terminal domain can form hydrophobic contacts with the NAC domain, while the polar and basic residues on the N-terminal domain interact with the acidic C-terminal domain. These interactions may protect the hydrophobic residues in the WT protein from interacting with other proteins before the N-terminal domain can interact with the lipid membrane.

We observed that elimination of either of the flanking domains led to major changes in the contacts formed between the remaining domains. When the N-terminal domain was removed, the overall number of NAC-C contacts increased (Figure 4a). The WT mostly sampled structures with few NAC-C contacts (Figure 3a), but the ΔN variant adopted structures with more NAC-C contacts than the WT variant. This was also observed qualitatively in the representative structures (Figure 2b). Interestingly, removal of the C-terminal domain led to a slight reduction in the number of contacts between the N-terminal and NAC domains (Figure 4b). Again, this can be seen qualitatively in the cluster representative structures (Figure 2C), where the N-terminal domain appears to be less tightly bound to the NAC domain.

**Figure 4.**
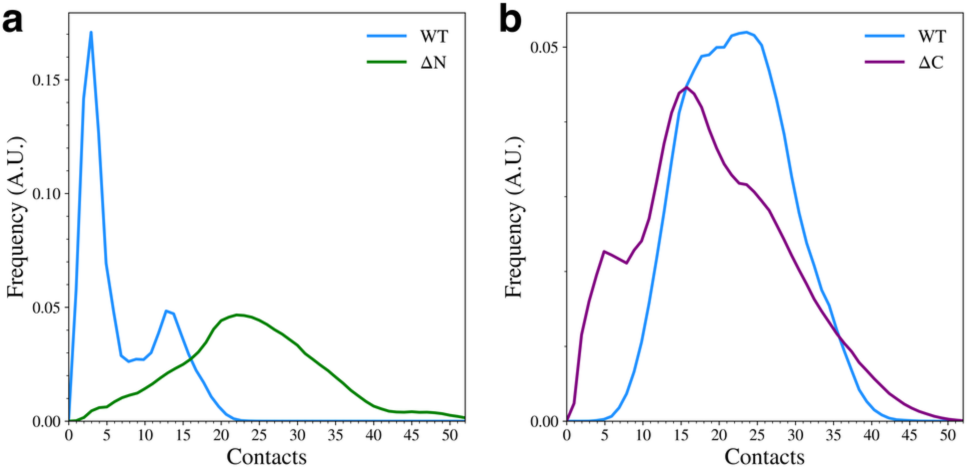
The distribution of NAC-C contacts (a) and NAC-N contacts (b) for the WT and domain deletion variants.

We also determined how the inter-domain hydrophobic contacts, hydrogen bonds, and salt bridges were affected by the removal of the flanking domains. Details of this contact analysis can be found in the Methods section. We found that the when the N-terminal domain was removed, the NAC-C hydrophobic contacts increased (Figure 5a), but when the C-terminal domain is removed, NAC-N hydrophobic contacts decreased (Figure 5b), with most of the sampled structures having no hydrophobic contacts between the two domains. The formation of hydrophobic NAC-N contacts observed in the WT protein may be dependent on N-C contacts, likely due to the amphipathic nature of the N-terminal domain and its ability to form contacts with the both the NAC and C-terminal domain. Removal of the N-terminal domain also increased the formation of NAC-C hydrogen bonds (Figure 5c), but removal of the C-terminal domain had little effect on the formation of NAC-N hydrogen bonds (Figure 5d). Lastly, the formation of NAC-C salt bridges (Figure 5e) or NAC-N salt bridges (Figure 5f) was almost unaffected by flanking domain removal. This is not surprising since significant salt bridge formation was mainly observed between the N-terminal and C-terminal domains.

**Figure 5.**
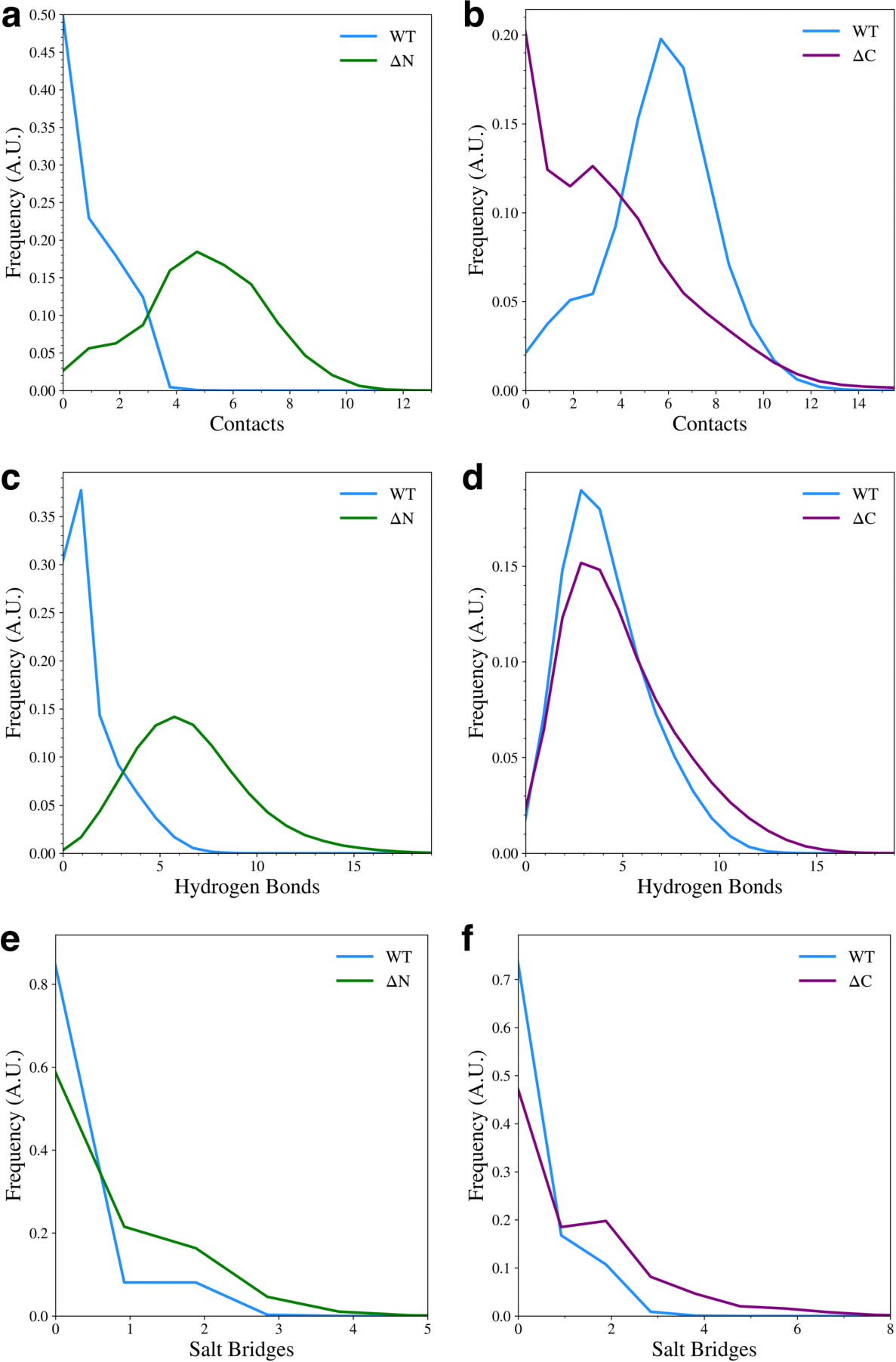
Contact distributions between the NAC and flanking domains for the WT and domain deletion variants. The plots show hydrophobic contacts between NAC-C (a) and NAC-N (b), hydrogen bonds between NAC-C (c) and NAC-N (d), and salt bridges between NAC-C (e) and NAC-N (f).

We also calculated the pairwise distances between C_α_ atoms for each pair of residues and used the most probable distance make a contact map (Figure 6). We only plotted distance values below 12 Å in our maps to aid in visualization. In Figure 6, the WT protein is shown along with either the ΔN (Figure 6a) or ΔC (Figure 6b) variants overlayed in inverted colors over the map. The individual contact maps for the ΔN (Figure S2a), and ΔC variants (Figure S2b) are also shown in Figure S2.

**Figure 6.**
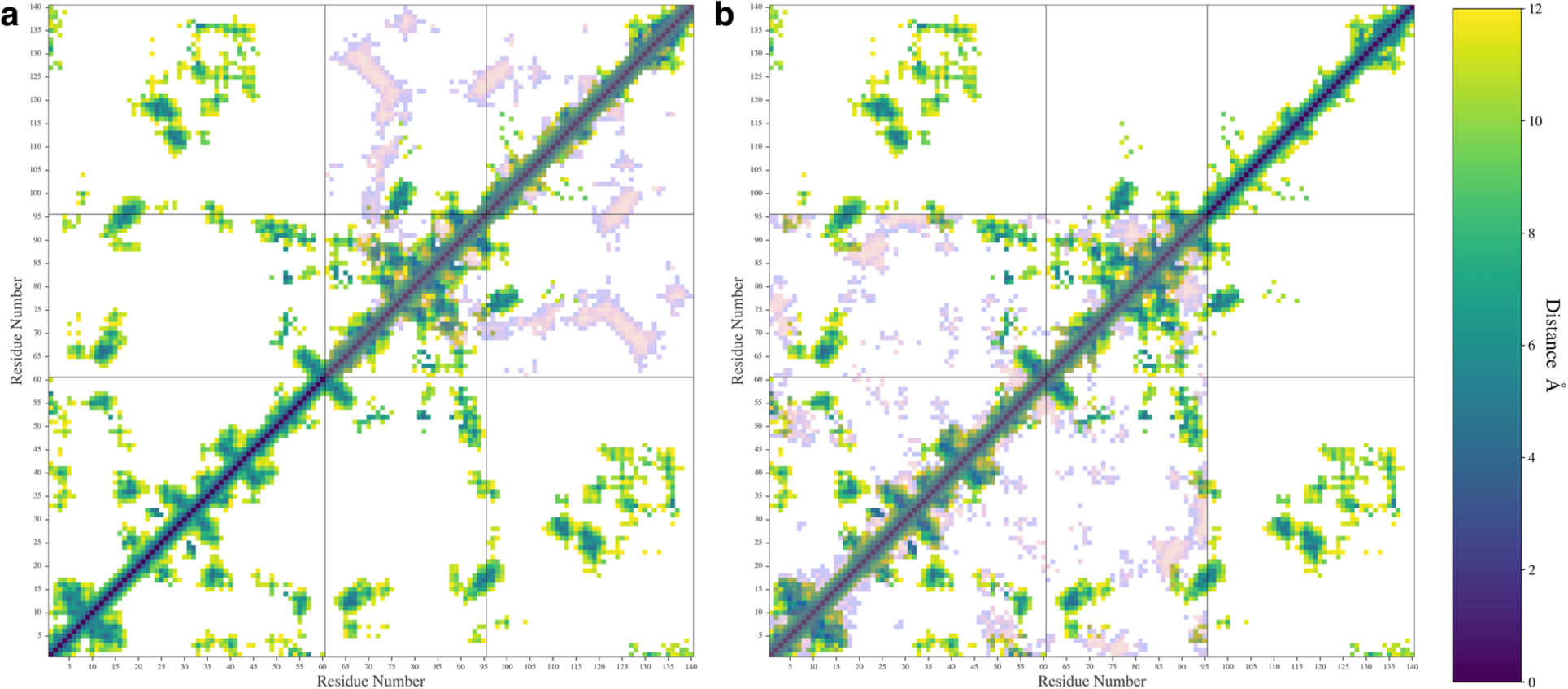
The contact maps showing the most probable distances between amino acid pairs for the WT protein with the ΔN (**a**) and ΔC (**b**) distances shown overlayed in inverted colors.

The contact map shows that the WT protein formed contacts between the N- and C-terminal domains and formed contacts between the NAC and N-terminal domains. The map also shows some contacts formed between the NAC and C-terminal domains, namely between the residues at the start of the C-terminal domain sequence and residues in the middle of the NAC domain sequence. This agrees with experimental results suggesting that the C-terminal domain can double back and form protective contacts with the NAC domain[19,36]. In the WT protein map, residues 20-45 on the N-terminal seem to be the most likely to have formed contacts with the C-terminal domain. Other N-terminal residues formed contacts with the NAC domain. The contact map shows close contacts between A27-I112, A27-L113, and A29-L113, which are all near A30. The A30P mutation is a well-studied mutation, though its effect on oligomerization is unclear[44]. NMR studies have shown that this mutation changes contacts between the N-terminal domain and the C-terminal domain[37]. Mutating the alanine to a proline is likely to reduce the flexibility of the N-terminal domain in a way that would disrupt the nearby N-C contacts that we observed. Additionally, the contact map shows an A53-V82 contact. The A53T mutation is also well studied and known to increase to propensity of the protein to form fibrils[45]. NMR studies have found that the A53T mutation decreases the flexibility of the N-terminal domain[37]. Mutating this contact could also possibly disrupt an interactions that could help hold the N-terminal domain in place to protect hydrophobic regions in the NAC domain.

The contact map for the WT also shows contacts formed between K45-D121 and K45-E123. The E46K mutation has been associated with an increase in aggregation and fibril formation[2,5]. WT fibrils have been found to form an E46-K80 salt bridge and the mutation of E46 to K46 leads to a polymorphic fibril that is more stable than the WT fibril[2]. Our observed interactions between K45-D121 and K45-E123 may be a contact that can help avoid the formation of the E46-K80 salt bridge that leads to WT fibrils.

The removal of the N-terminal domain (Figure 6a, S2a) shows contacts that indicate the presence of a Δ-sheet made up of residues 68-74 on the NAC domain and residues 123-130 on the C-terminal domain. This β-sheet motif includes many hydrophobic residues, specifically, A69, V70, V71, V74, A124, Y125, M127, and P128. Removal of the C-terminal domain (Figure 6b, S2b) resulted in a change in the contacts between the NAC and N-terminal domain, especially in the first 10 residues of the protein and in the 20-45 region that formed contacts with the C-terminal domain in the WT protein. Removal of the domain also led to an increase in the intra-domain contacts formed in the N-terminal and NAC domains. This may explain our previous observation that removal of the domain does not lead to a significant increase in NAC-N contacts. It appears that both N-terminal and NAC domains begin to self-interact more when the C-terminal domain is removed. Removal of the C-terminal domain also seems to disrupt some of the β-sheet motifs observed in the WT type contact map.

### Flanking Domain Removal Affects Secondary Structure Formation

We analyzed the presence of secondary structure motifs to determine how the removal of the flanking domains affected the propensity to form β-sheets. The average number of residues adopting either an α-helix or β-sheet motif was determined using DSSP [46]. For the entire protein (Figure 7a), we found that the removal of the N-terminal domain led to an increase in Δ-sheet content, while removal of the C-terminal domain decreased Δ-sheet content with respect to the WT protein. We also observed that the isoNAC variant sampled less α-helical or Δ-sheet content overall. Focusing only on the NAC domain (Figure 7b), the ΔN variant showed an increase in Δ-sheet content. Interestingly it also showed a slight increase in sampled helicity. In the N-terminal domain (Figure 7c), we found that the removal of the C-terminal domain appeared to disrupt the formation of β-sheet content, which was also observed in the contact maps. Lastly, we observed that removing the N-terminal domain led to a significant increase in C-terminal domain Δ-content, compared to the WT protein (Figure 7d), which had very little Δ-content in the C-terminal domain. Comparing these results with the representative cluster structures (Figures 1 and 2) and the contact maps (Figure 6), they suggest that N-terminal domain contacts between the NAC and C-terminal domains may lower the propensity to form inter-domain β-sheets. When the N-terminal domain is removed, the NAC and C-terminal domains may sample more Δ-motifs than the WT protein.

**Figure 7.**
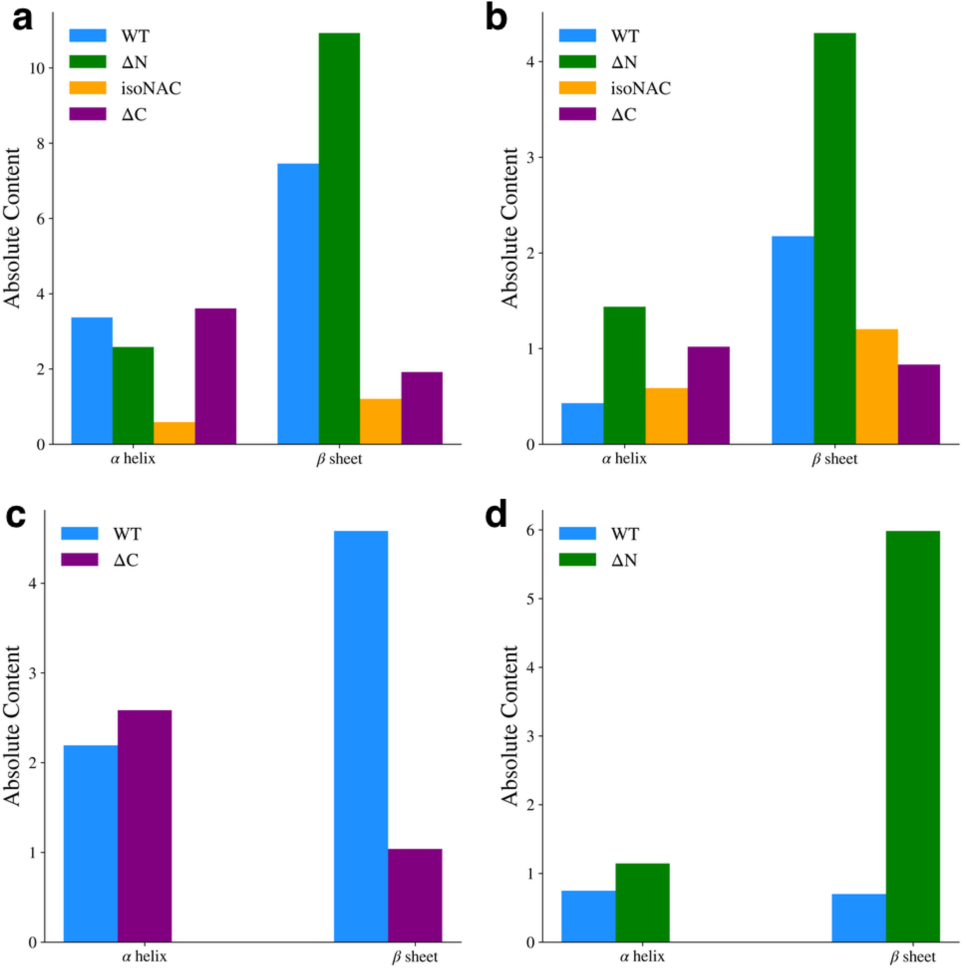
Average α-helix and β-sheet content for the full protein (a), the NAC domain (b) the N-terminal domain (c), and the C-terminal domain (d).

### Removal of Flanking Domains Increases Protein Compactness

We found that removal of either flanking domain led to an increase in compactness for the individual domains, compared to same domain in the WT protein. The compactness was measured by calculating the radius of gyration, *Rg* for each individual domain. We also calculated the distribution of *Rg* for the full proteins (Figure S3), which generally followed the expected size-dependent differences for the WT and isoNAC variants and did not give much meaningful insight into the effect of removing the flanking domains. Although the ΔN and ΔC variants both have a slightly different number of residues (80 and 95, respectively), they showed similar *Rg* distributions, suggesting the domain-deletion mutant proteins, as a whole, show similar levels of compactness.

Removal of the flanking domains led to an increase in compactness of the N-terminal, NAC, and C-terminal domains (Figure 8). Removal of the C-terminal domain (Figure 8A) increased the sampling of structures with a more compact N-terminal domain. The compactness of the NAC domain also increased when both flanking domains were removed (Figure 8b). The isoNAC variant showed increased compactness compared to the WT and domain deletion variants. Removal of either the N-terminal or C-terminal domains, individually, also increased the compactness of the NAC domain when compared to the WT protein, though not as much as the isoNAC variant. The distributions of the NAC domain *Rg* values for both the ΔN and ΔC variants were similar, which suggests that removing either flanking domain may result in a comparable increase in compactness of the NAC domain. Removal of the N-terminal domain, however, lead to a larger increase in the compactness of the C-terminal domain (Figure 8c). This coincides with the observed increase in β-sheet motifs sampled by the ΔN variant.

**Figure 8.**
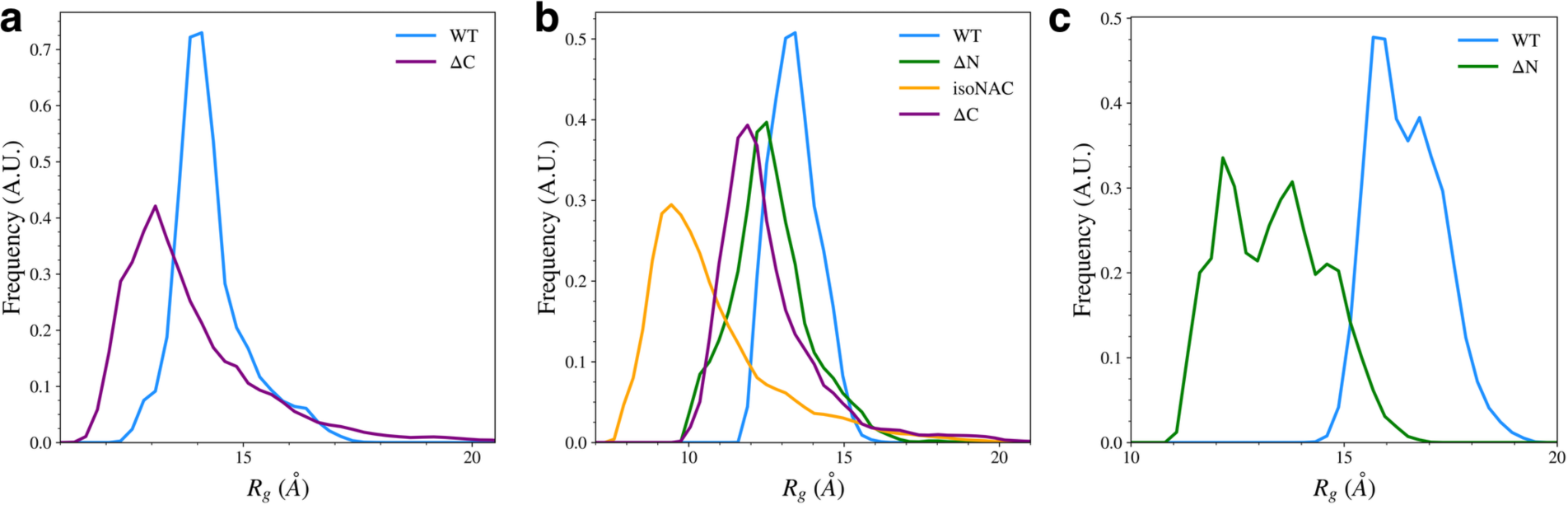
The radius of gyration, *Rg*, for the N-terminal domain (a), NAC domain (b), and C-terminal domains (c).

Overall, the *Rg* distributions suggest that one of the roles of the two flanking domains is to maintain the elongation of the protein. Removal of either the N-terminal or C-terminal domain led to an increase in compactness for the other flanking domain. This effect was especially prominent when the N-terminal domain was removed. Removal of either domain also seems to increase the compactness of the NAC domain. This suggests that both flanking domains may be protective against aggregation by increasing elongation in the NAC domain. Because the N-terminal domain can form contacts with both the C-terminal and NAC domain, it is possible that the N-C contacts may help maintain elongation in the flanking domains which is then imposed via NAC-N contacts on the NAC domain. Either NAC-N or NAC-C interactions alone appear to maintain some elongation in the NAC domain, but having both flanking domains on the protein together reduces the compactness more.

We also measured the total solvent accessible surface area (SASA) for the domains in each variant (Figure 9, top) and the SASA of the hydrophobic residues (Figure 9, bottom) to understand how the flanking domains may modulate the exposure of hydrophobic residues to the solvent. The hydrophobic SASA is likely to give some insight into how protein-protein interactions between monomers may occur in the cell. Although our simulations only consider the behavior of protein monomers, a conformation with a large amount of exposed hydrophobic surface area is likely be more prone to formation of toxic aggregates.

**Figure 9.**
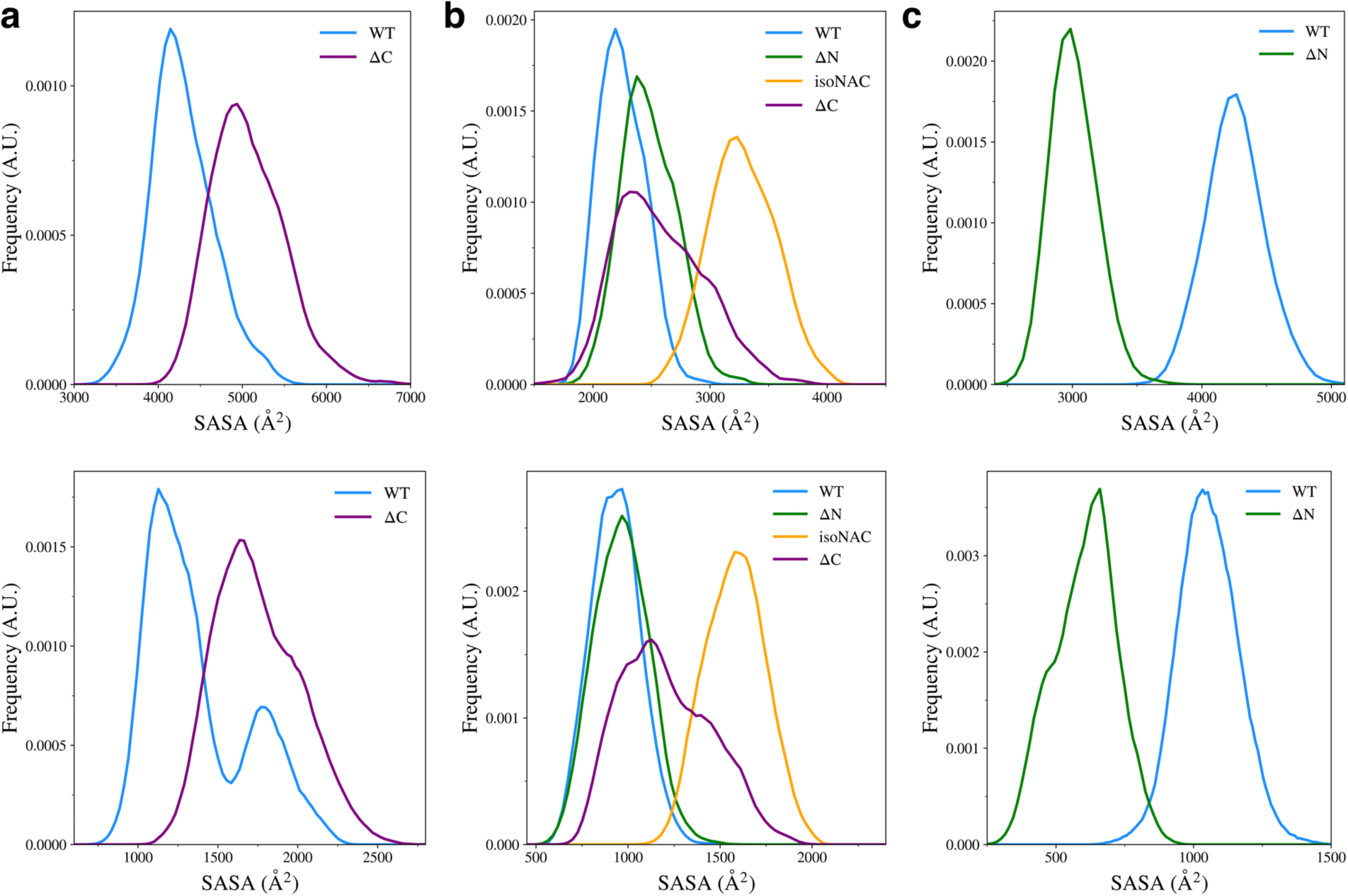
Solvent accessible surface area of all residues (top) and the hydrophobic residues (bottom) for the N-terminal domain (a), the NAC domain (b), and the C-terminal domain (c).

We found that removal of the C-terminal domain led to an increase in conformations with higher N-terminal domain SASA (figure 9a, top) and increased exposed hydrophobic surface area (figure 9a, bottom). Experimental work has shown that removal of the C-terminal domain leads to aggregation of protein monomers[24,30,47]. Our work suggests that a possible mechanism for this may be that when the C-terminal domain is removed, the N-terminal domain is more likely to adopt structures that have increased compactness with more exposed hydrophobic residues, compared to the WT asyn.

We also found that when the C-terminal domain is removed, compared to the WT, the ΔC variant showed an increase in the NAC domain SASA (Figure 9b, top) and an increase in the exposed hydrophobic surface area (Figure 9b, bottom). Interestingly, removal of the N-terminal domain led to a slight increase in SASA and almost no change in the surface area of hydrophobic residues. The removal of both flanking domains caused the NAC to have increased SASA and increased hydrophobic surface area. Without either flanking domain forming contacts with the NAC domain, it seems that most of the sampled structures had increased hydrophobic exposure to the solution. The ΔN variant also showed a large decrease in overall and hydrophobic SASA of the C-terminal domain (Figure 9C). It appears that increase in contacts between the NAC and C-terminal domains also keeps the hydrophobic residues from the solvent.

### Interdomain Interactions Modulate the Protein Electrostatic Potential

An important role of the N-terminal and NAC domains is their interaction with lipid membranes to fold the N-terminal domain in the membrane[1,11,13-15,17,48,49]. The N-terminal domain is amphipathic and can form a helix that folds peripherally into the lipid membrane, along with part of the NAC domain[15,32,48]. When this occurs, the C-terminal domain then is projected into the cytosol, where it can act as a scaffold to interact with other proteins[17]. Part of this folding process involves interactions between the protein and negatively charged lipid headgroups on the lipid membrane surface. Additionally, electrostatic interactions between monomers can play an important role in the formation of asyn fibrils[2,26,32,50]. To understand how the removal of either flanking domain might affect electrostatic interactions with the asyn monomer, we calculated the electrostatic potential (ESP) of the representative structures of the most populated clusters.

We calculated the ESP using VMD[51] and mapped the values to the SASA of the protein, using a voltage range of −5*k*_*B*_*T*/*e* to −5*k*_*B*_*T*/*e*, where *k*_*B*_ is Boltzmann’s constant, *T* is the temperature (310 K) and *e* is the charge of an electron (Figure 10, top). We also visualized these same structures using the SASA representation colored to show the specific domains (Figure 10, bottom). We included the domain coloring to show how the ESP manifests on the individual domains. We found that the WT protein structure (Figure 10A) showed localized areas of positive ESP on the N-terminal and NAC domains that correspond to the locations of imperfect KTKEGV repeat units. The bottom left hand blue area is made up of the KTKEGV motif beginning with residue 58, the bottom right-hand region of positive potential is the KTVEGA motif beginning at residue 80, and the top right hand positive region is the KTKQGV repeat beginning at residue 21. This conformation can orient itself in such a way that the N-terminal and C-terminal domains can interact with negatively charged lipid headgroups and the C-terminal domain can be oriented toward the cytosol. This may aid the WT protein in selectively binding to negatively charged membrane surfaces and may also aid the folding of the N-terminal domains into the cell membrane by creating electrostatic anchor points on the lipid membrane. Overall, this type of interaction may lower the barrier required to fold the N-terminal domain into the membrane by keeping it close to anionic lipid headgroups. While we believe this is an intriguing suggestion that should be further investigated, verifying this will require further simulations of the protein structures with charged and uncharged lipid membranes.

**Figure 10.**
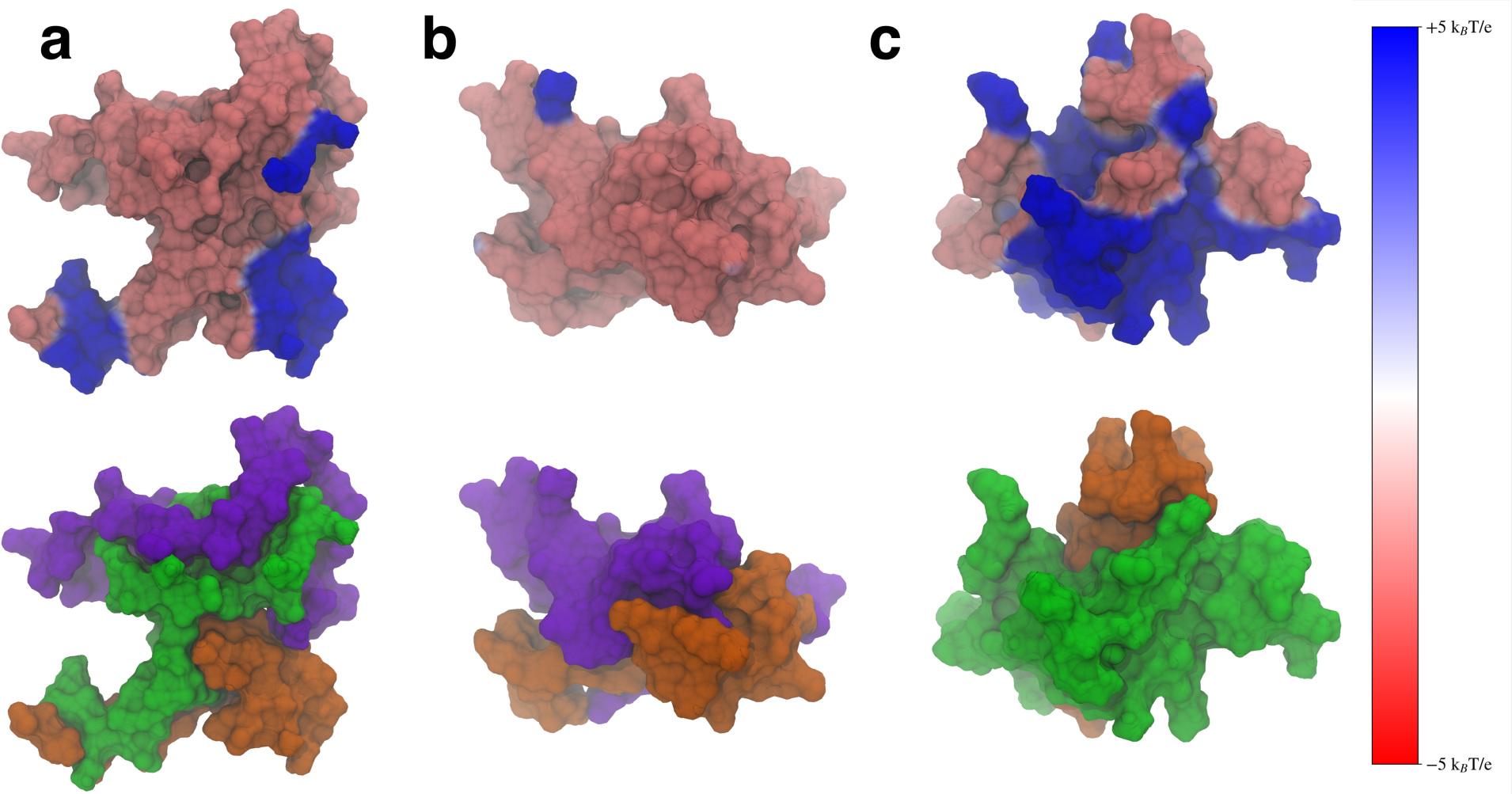
The electrostatic potential mapped onto the surface representation (top) and the surface representation colored by domain (bottom) for the WT (a), ΔN (b), and ΔC (c) variants. The N-terminal domain is shown in green, the NAC in orange, and the C-terminal domain in violet. The positive electrostatic potential is shown in blue and the negative potential in red.

When the N-terminal domain is removed, the ESP of the structure does not show the distinct regions of positive ESP seen in the WT (Figure 10b). The structure shows mostly negative ESP, which is likely to impair the ability of the protein to interact with lipid membranes that contain negatively charged lipid headgroups. Removal of the C-terminal domain does lead to a large increase of positive ESP on the surface of the molecule (Figure 10C). This increased positive surface is larger than the WT and could lead to stronger electrostatic interactions between the protein and the membrane. Experimental work has found that C-terminal truncations lead to increased protein-membrane interaction[30], which may be due to this increase in positive ESP on the surface of the protein. When the C-terminal truncated protein encounters a lipid membrane, the stronger interactions may also make it harder for the protein to fold into the lipid membrane, having to overcome a larger barrier when dissociating from the charged lipid headgroups to fold the N-terminal domain into the membrane. We believe these results suggest that both flanking domains can help modulate the ESP of the protein so that the WT protein can balance its interactions with negatively charged lipids in a way that does not bind too strongly to the membrane. Disruption of the interdomain interactions, such as through a domain deletion, leads to structures that may not be able to bind to negative lipids or lead to over-stabilized protein-membrane contacts.

Our observed electrostatic potentials may also have implications for the role of the flanking domains in protecting against protein aggregation. C-terminal truncations of asyn have been shown to increase the strength of protein-protein interactions[30]. Large regions of positive electrostatic potential, such as those seen in the ΔC variant would likely help over-stabilize protein-protein interactions compared to the WT asyn. This could cause the formation of protein structures that are more difficult to disassemble when needed, leading to the formation of toxic aggregates.

### A Model for Flanking-Domain Cooperation

Our results suggest that the protective role of the flanking domains may be due to cooperation between the N and C -terminal domains. The C-terminal domain is the only domain in asyn that contains proline residues. The five prolines in this domain help the C-terminal domain remain elongated. The formation of N-C contacts may allow the C-terminal domain to impose this elongation on the N-terminal domain. The NAC-N contacts may then also help maintain some elongation in the NAC domain, possibly protecting against compact and toxic NAC conformations. We found that when the N-terminal domain was removed, the direct contacts between the NAC and C-terminal domains led to an increase in β-rich conformations, along with an increase in compactness. Additionally, an increase in structures with a more compact NAC domain was also observed. Removal of the C-terminal domain also resulted in increased compactness in the NAC and N-terminal domains. The C-terminal domain deletion also led to structures that showed an increase in hydrophobic SASA in both the NAC and N-terminal domains. We believe it is possible that both flanking domains work together to help the protein sample structures that both maintain elongation of the NAC domain and protect hydrophobic residues from being exposed to the solvent.

We also found that the contacts between the N- and C-terminal domains can modulate the surface ESP of the WT such that the positive ESP is focused on the surface KTKEGV imperfect repeat units. A key role of the N-terminal domain is to form helical structures within the lipid membrane. The ESP surface we found for the WT protein may help the protein to bind to negative lipid surfaces without over stabilizing protein-membrane interactions. We are proposing that this may allow the protein to remain on the membrane surface while inserting hydrophobic residues into the membrane for folding, without increasing the energy barrier for protein to unbind from the membrane lipids. It should be noted, however, that we would need to run protein-lipid binding simulations to test this hypothesis, which we plan to investigate in future work.

## Conclusion

In this work we used GAMD simulations to model the effects of removing the N-terminal or C-terminal domains of asyn monomers, so that we could observe what role they play in modulating the structure of the protein. We found that the two flanking domains may work cooperatively to maintain the elongation of all three domains. Because of its amphipathic character, the N-terminal domain can form salt-bridges and hydrogen bonds with the C-terminal domain, and hydrophobic interactions, along with hydrogen bonds, with the NAC domain. Removal of the N-terminal domain leads to increased interactions between the NAC and C-terminal domains that result in increased sampling of β-sheet motifs. Removal of the C-terminal domain does not seem to lead to a significant increase in contacts or Δ-motifs but does seem to disrupt the specificity of the interactions formed between the NAC and C-terminal domains.

We observed that removal of either domain led to an increase in compactness in the remaining domains. This was notable in the C-terminal domain, which became more compact when the N-terminal domain was removed, likely due to the increased sampling in β-sheet structures between the NAC and C-terminal domain. Interestingly, we found that removal of the C-terminal domain led to an increase in the amount of hydrophobic SASA of the protein. Removal of the N-terminal domain did not increase the hydrophobic SASA in the NAC domain and reduced the amount of SASA in the C-terminal domain. This may suggest that N-terminal deletions do not form aggregates initially via monomer-monomer hydrophobic interactions, however, this is a result that requires further investigation to fully understand its effect in asyn function.

We also observed that the WT asyn can adopt a structure with an ESP that could possibly selectively bind to negatively charged lipid membranes. This observed structure would orient the N-terminal and NAC domains towards the membrane and the C-terminal domain away from the membrane. This type of structure could also hold the protein in place, possibly reducing the barrier to fold the N-terminal domain into the membrane, however, further simulations would be required to test this proposed hypothesis. Removing the N-terminal domain led to a structure that had almost no positive charge, but removing the C-terminal domain led to an increase in the positive ESP. We expect that although this type of structure would interact with negatively charged lipids, it may also increase the barrier required to fold the N-terminal domain into the lipid membrane, due to the increased electrostatic interactions between the protein and the membrane. Additionally, this increase in positive ESP could lead to stronger protein-protein interactions that may result in the formation of structures that are overly stable and could lead to toxic aggregates; however, as said above, more research is required to explore this further.

We believe these results suggest that both domains work cooperatively to both protect against toxic aggregation and to modulate the ESP so that the protein can interact favorably with negative lipids in the membrane. When the N-terminal domain binds to both the C-terminal and NAC domains, the flanking domains form electrostatic interactions that could keep domains elongated. The hydrophobic NAC-N interactions then may also help maintain elongation in the NAC domain. This may also protect hydrophobic residues on the NAC domain from the solvent. The resulting structures that are sampled by the protein appear to maintain an ESP that helps the N-terminal domain fold into the membrane and does not lead to over-stabilized protein-membrane or protein-protein interactions.

## Methods

Four variants were simulated in this study: A WT asyn protein (residues 1-140), and three domain deletion variants: ΔN (residues 61-140), ΔC (residues 1-95), and isoNAC (residues 61-95). MD simulations were performed using the Amber 16 software package[52] and the CHARMM36m force field[53]. A time step of 4 fs using hydrogen mass repartitioning[54] was employed for all simulations.

The MD simulation setup was prepared using the following procedure. A fully extended structure was constructed from the protein sequence using the Molefacture plugin in VMD[55]. The proteins were energy minimized and collapsed *in vacuo* using MD at 300 K. The collapsed proteins were then solvated with a 20 Å or 25 Å layer of TIP3P water and neutralized with NaCl to a concentration of 0.15 M. The resulting WT protein had 128,920 atoms, the N-terminus deletion variant had 77,397 atoms, the NAC domain had 34,071 atoms, and the C-terminus deletion variant had 39,380 atoms.

The energy was then minimized, and the solvent density was relaxed under constant pressure and temperature. After this, the volume was held constant, and the system was heated from 310 K to 600 K and equilibrated at high temperature for 300 ns. This reverse annealing was used to ensure that we sampled fully unfolded structures during the enhanced sampling simulations. During the heating stage, the trajectories were visually inspected in VMD to ensure that proteins were not self-interacting with periodic images.

Because α-synuclein is an IDP, we used Gaussian accelerated molecular dynamics (GAMD) to enhance the sampling of our simulations[39,40]. We did not use the reweighting procedure. The GAMD parameters were chosen using the standard advice in the Amber16 manual. Simulations were run for at least 0.6 μs with GAMD at 310 K for all systems. Coordinates were recorded every 1 ps and the first 150 ns of each trajectory were discarded for analysis. Analyses on the trajectories were performed using Pytraj[56], NumPy[57], and SciPy[58] libraries. All plots were made using the Matplotlib library[59].The clustering analysis was done using the k-means method[43]. Contact between residues was defined as having a distance between C_α_ atoms less than 6 Å. Hydrophobic contacts were defined the same way between residues with hydrophobic sidechains. Hydrogen bonds were defined using the default Pytraj definition, with a distance cutoff of 3.0 Å and an angle cutoff of 135°. Salt bridge contacts were defined as oppositely charged residues where the distance between an N or O atom was within 4 Å. Secondary structure analyses were conducted using the Dictionary of Secondary Structure of Proteins (DSSP)[46] with the MDTraj library[46,60]. The ESP calculations were done using the PMEplot plugin in VMD[51]. VMD was used to visualize and render all molecular structures[55].

## Acknowledgments

This work was run on hardware resources supported by the National Institutes of Health under award number NIH SC2GM131992 and NSF award number 2320718.

## REFERENCES

1. Auluck, P.K.; Caraveo, G.; Lindquist, S. alpha-Synuclein: Membrane Interactions and Toxicity in Parkinson’s Disease. In Annual Review of Cell and Developmental Biology, Vol 26, Schekman, R., Goldstein, L., Lehmann, R., Eds.; Annual Review of Cell and Developmental Biology; Annual Reviews: Palo Alto, 2010; Volume 26, pp. 211-233.

2. Boyer, D.R.; Li, B.; Sun, C.; Fan, W.; Zhou, K.; Hughes, M.P.; Sawaya, M.R.; Jiang, L.; Eisenberg, D.S. The α-synuclein hereditary mutation E46K unlocks a more stable, pathogenic fibril structure. Proceedings of the National Academy of Sciences 2020, 117, 3592–3602, doi:doi:10.1073/pnas.1917914117.

3. Meade, R.M.; Fairlie, D.P.; Mason, J.M. Alpha-synuclein structure and Parkinson’s disease – lessons and emerging principles. Molecular Neurodegeneration 2019, 14, 29, doi:10.1186/s13024-019-0329-1.

4. Tuttle, M.D.; Comellas, G.; Nieuwkoop, A.J.; Covell, D.J.; Berthold, D.A.; Kloepper, K.D.; Courtney, J.M.; Kim, J.K.; Barclay, A.M.; Kendall, A.;, et al. Solid-state NMR structure of a pathogenic fibril of full-length human alpha-synuclein. Nature structural & molecular biology 2016, 23, 409–415, doi:10.1038/nsmb.3194.

5. Zarranz, J.J.; Alegre, J.; Gomez-Esteban, J.C.; Lezcano, E.; Ros, R.; Ampuero, I.; Vidal, L.; Hoenicka, J.; Rodriguez, O.; Atares, B.;, et al. The new mutation, E46K, of alpha-synuclein causes Parkinson and Lewy body dementia. Annals of neurology 2004, 55, 164–173, doi:10.1002/ana.10795.

6. Lashuel, H.A.; Overk, C.R.; Oueslati, A.; Masliah, E. The many faces of α-synuclein: from structure and toxicity to therapeutic target. Nature Reviews Neuroscience 2012, 14, 38, doi:10.1038/nrn3406.

7. Abeliovich, A.; Schmitz, Y.; Fariñas, I.; Choi-Lundberg, D.; Ho, W.-H.; Castillo, P.E.; Shinsky, N.; Verdugo, J.M.G.; Armanini, M.; Ryan, A.;, et al. Mice Lacking &#x3b1;-Synuclein Display Functional Deficits in the Nigrostriatal Dopamine System. Neuron 2000, 25, 239–252, doi:10.1016/S0896-6273(00)80886-7.

8. Cabin, D.E.; Shimazu, K.; Murphy, D.; Cole, N.B.; Gottschalk, W.; McIlwain, K.L.; Orrison, B.; Chen, A.; Ellis, C.E.; Paylor, R.;, et al. Synaptic Vesicle Depletion Correlates with Attenuated Synaptic Responses to Prolonged Repetitive Stimulation in Mice Lacking α-Synuclein. The Journal of Neuroscience 2002, 22, 8797–8807, doi:10.1523/jneurosci.22-20-08797.2002.

9. Murphy, D.D.; Rueter, S.M.; Trojanowski, J.Q.; Lee, V.M.-Y. Synucleins Are Developmentally Expressed, and α-Synuclein Regulates the Size of the Presynaptic Vesicular Pool in Primary Hippocampal Neurons. The Journal of Neuroscience 2000, 20, 3214–3220, doi:10.1523/jneurosci.20-09-03214.2000.

10. Nemani, V.M.; Lu, W.; Berge, V.; Nakamura, K.; Onoa, B.; Lee, M.K.; Chaudhry, F.A.; Nicoll, R.A.; Edwards, R.H. Increased Expression of &#x3b1;-Synuclein Reduces Neurotransmitter Release by Inhibiting Synaptic Vesicle Reclustering after Endocytosis. Neuron 2010, 65, 66–79, doi:10.1016/j.neuron.2009.12.023.

11. Kamp, F.; Beyer, K. Binding of alpha-synuclein affects the lipid packing in bilayers of small vesicles. J. Biol. Chem. 2006, 281, 9251–9259, doi:10.1074/jbc.M512292200.

12. Perlmutter, J.D.; Braun, A.R.; Sachs, J.N. Curvature Dynamics of alpha-Synuclein Familial Parkinson Disease Mutants MOLECULAR SIMULATIONS OF THE MICELLE-AND BILAYER-BOUND FORMS. J. Biol. Chem. 2009, 284, 7177–7189, doi:10.1074/jbc.M808895200.

13. Davidson, W.S.; Jonas, A.; Clayton, D.F.; George, J.M. Stabilization of alpha-synuclein secondary structure upon binding to synthetic membranes. J. Biol. Chem. 1998, 273, 9443–9449, doi:10.1074/jbc.273.16.9443.

14. Braun, A.R.; Sevcsik, E.; Chin, P.; Rhoades, E.; Tristram-Nagle, S.; Sachs, J.N. α-Synuclein Induces Both Positive Mean Curvature and Negative Gaussian Curvature in Membranes. Journal of the American Chemical Society 2012, 134, 2613–2620, doi:10.1021/ja208316h.

15. Braun, A.R.; Lacy, M.M.; Ducas, V.C.; Rhoades, E.; Sachs, J.N. α-Synuclein-Induced Membrane Remodeling Is Driven by Binding Affinity, Partition Depth, and Interleaflet Order Asymmetry. Journal of the American Chemical Society 2014, 136, 9962–9972, doi:10.1021/ja5016958.

16. Ullman, O.; Fisher, C.K.; Stultz, C.M. Explaining the Structural Plasticity of α-Synuclein. Journal of the American Chemical Society 2011, 133, 19536–19546, doi:10.1021/ja208657z.

17. Eliezer, D.; Kutluay, E.; Bussell, R.; Browne, G. Conformational properties of alpha-synuclein in its free and lipid-associated states. J. Mol. Biol. 2001, 307, 1061–1073, doi:10.1006/jmbi.2001.4538.

18. Uversky, V.N.; Li, J.; Fink, A.L. Evidence for a Partially Folded Intermediate in &#x3b1;-Synuclein Fibril Formation *. J. Biol. Chem. 2001, 276, 10737–10744, doi:10.1074/jbc.M010907200.

19. Bertoncini, C.W.; Jung, Y.-S.; Fernandez, C.O.; Hoyer, W.; Griesinger, C.; Jovin, T.M.; Zweckstetter, M. Release of long-range tertiary interactions potentiates aggregation of natively unstructured α-synuclein. Proceedings of the National Academy of Sciences 2005, 102, 1430–1435, doi:doi:10.1073/pnas.0407146102.

20. Theillet, F.-X.; Binolfi, A.; Bekei, B.; Martorana, A.; Rose, H.M.; Stuiver, M.; Verzini, S.; Lorenz, D.; van Rossum, M.; Goldfarb, D.;, et al. Structural disorder of monomeric α-synuclein persists in mammalian cells. Nature 2016, 530, 45–50, doi:10.1038/nature16531.

21. Kessler, J.C.; Rochet, J.-C.; Lansbury, P.T. The N-Terminal Repeat Domain of α-Synuclein Inhibits β-Sheet and Amyloid Fibril Formation. Biochemistry 2003, 42, 672–678, doi:10.1021/bi020429y.

22. Sorrentino, Z.A.; Giasson, B.I. The emerging role of &#x3b1;-synuclein truncation in aggregation and disease. J. Biol. Chem. 2020, 295, 10224–10244, doi:10.1074/jbc.REV120.011743.

23. Michell, A.W.; Tofaris, G.K.; Gossage, H.; Tyers, P.; Spillantini, M.G.; Barker, R.A. The Effect of Truncated Human &#945;-Synuclein (1&#8211;120) on Dopaminergic Cells in a Transgenic Mouse Model of Parkinson’s Disease. Cell Transplantation 2007, 16, 461–474, doi:10.3727/000000007783464911.

24. Iyer, A.; Roeters, S.J.; Kogan, V.; Woutersen, S.; Claessens, M.M.A.E.; Subramaniam, V. C-Terminal Truncated α-Synuclein Fibrils Contain Strongly Twisted β-Sheets. Journal of the American Chemical Society 2017, 139, 15392–15400, doi:10.1021/jacs.7b07403.

25. Killinger, B.A.; Madaj, Z.; Sikora, J.W.; Rey, N.; Haas, A.J.; Vepa, Y.; Lindqvist, D.; Chen, H.; Thomas, P.M.; Brundin, P.;, et al. The vermiform appendix impacts the risk of developing Parkinson’s disease. Science Translational Medicine 2018, 10, eaar5280, doi:10.1126/scitranslmed.aar5280.

26. van der Wateren, I.M.; Knowles, T.P.J.; Buell, A.K.; Dobson, C.M.; Galvagnion, C. C-terminal truncation of α-synuclein promotes amyloid fibril amplification at physiological pH. Chemical Science 2018, 9, 5506–5516, doi:10.1039/C8SC01109E.

27. Ni, X.; McGlinchey, R.P.; Jiang, J.; Lee, J.C. Structural Insights into α-Synuclein Fibril Polymorphism: Effects of Parkinson’s Disease-Related C-Terminal Truncations. J. Mol. Biol. 2019, 431, 3913–3919, 10.1016/j.jmb.2019.07.001.

28. Gallardo, J.; Escalona-Noguero, C.; Sot, B. Role of α-Synuclein Regions in Nucleation and Elongation of Amyloid Fiber Assembly. ACS chemical neuroscience 2020, 11, 872–879, doi:10.1021/acschemneuro.9b00527.

29. Hass, E.W.; Sorrentino, Z.A.; Xia, Y.; Lloyd, G.M.; Trojanowski, J.Q.; Prokop, S.; Giasson, B.I. Disease-, region- and cell type specific diversity of α-synuclein carboxy terminal truncations in synucleinopathies. Acta Neuropathologica Communications 2021, 9, 146, doi:10.1186/s40478-021-01242-2.

30. Zhang, C.; Pei, Y.; Zhang, Z.; Xu, L.; Liu, X.; Jiang, L.; Pielak, G.J.; Zhou, X.; Liu, M.; Li, C. C-terminal truncation modulates α-Synuclein’s cytotoxicity and aggregation by promoting the interactions with membrane and chaperone. Communications Biology 2022, 5, 798, doi:10.1038/s42003-022-03768-0.

31. Röntgen, A.; Toprakcioglu, Z.; Tomkins, J.E.; Vendruscolo, M. Modulation of α-synuclein in vitro aggregation kinetics by its alternative splice isoforms. Proceedings of the National Academy of Sciences 2024, 121, e2313465121, doi:doi:10.1073/pnas.2313465121.

32. Lorenzen, N.; Lemminger, L.; Pedersen, J.N.; Nielsen, S.B.; Otzen, D.E. The N-terminus of α-synuclein is essential for both monomeric and oligomeric interactions with membranes. FEBS Letters 2014, 588, 497–502, 10.1016/j.febslet.2013.12.015.

33. McGlinchey, R.P.; Ni, X.; Shadish, J.A.; Jiang, J.; Lee, J.C. The N terminus of α-synuclein dictates fibril formation. Proceedings of the National Academy of Sciences 2021, 118, e2023487118, doi:doi:10.1073/pnas.2023487118.

34. Prasad, K.; Beach Thomas, G.; Hedreen, J.; Richfield Eric, K. Critical Role of Truncated α-Synuclein and Aggregates in Parkinson’s Disease and Incidental Lewy Body Disease. Brain Pathology 2012, 22, 811–825, doi:10.1111/j.1750-3639.2012.00597.x.

35. Bernadó, P.; Bertoncini, C.W.; Griesinger, C.; Zweckstetter, M.; Blackledge, M. Defining Long-Range Order and Local Disorder in Native α-Synuclein Using Residual Dipolar Couplings. Journal of the American Chemical Society 2005, 127, 17968–17969, doi:10.1021/ja055538p.

36. Dedmon, M.M.; Lindorff-Larsen, K.; Christodoulou, J.; Vendruscolo, M.; Dobson, C.M. Mapping Long-Range Interactions in α-Synuclein using Spin-Label NMR and Ensemble Molecular Dynamics Simulations. Journal of the American Chemical Society 2005, 127, 476–477, doi:10.1021/ja044834j.

37. Okuwaki, R.; Shinmura, I.; Morita, S.; Matsugami, A.; Hayashi, F.; Goto, Y.; Nishimura, C. Distinct residual and disordered structures of alpha-synuclein analyzed by amide-proton exchange and NMR signal intensity. Biochimica et Biophysica Acta (BBA) - Proteins and Proteomics 2020, 1868, 140464, 10.1016/j.bbapap.2020.140464.

38. Kumari, P.; Ghosh, D.; Vanas, A.; Fleischmann, Y.; Wiegand, T.; Jeschke, G.; Riek, R.; Eichmann, C. Structural insights into α-synuclein monomer–fibril interactions. Proceedings of the National Academy of Sciences 2021, 118, e2012171118, doi:doi:10.1073/pnas.2012171118.

39. Miao, Y.; Sinko, W.; Pierce, L.; Bucher, D.; Walker, R.C.; McCammon, J.A. Improved Reweighting of Accelerated Molecular Dynamics Simulations for Free Energy Calculation. J. Chem. Theory Comput. 2014, 10, 2677–2689, doi:10.1021/ct500090q.

40. Miao, Y.; Feher, V.A.; McCammon, J.A. Gaussian Accelerated Molecular Dynamics: Unconstrained Enhanced Sampling and Free Energy Calculation. J. Chem. Theory Comput. 2015, 11, 3584–3595, doi:10.1021/acs.jctc.5b00436.

41. Hamelberg, D.; Mongan, J.; McCammon, J.A. Accelerated molecular dynamics: A promising and efficient simulation method for biomolecules. The Journal of Chemical Physics 2004, 120, 11919–11929, doi:10.1063/1.1755656.

42. Hamelberg, D.; de Oliveira, C.A.F.; McCammon, J.A. Sampling of slow diffusive conformational transitions with accelerated molecular dynamics. The Journal of Chemical Physics 2007, 127, 155102, doi:10.1063/1.2789432.

43. Shao, J.; Tanner, S.W.; Thompson, N.; Cheatham, T.E. Clustering Molecular Dynamics Trajectories: 1. Characterizing the Performance of Different Clustering Algorithms. J. Chem. Theory Comput. 2007, 3, 2312–2334, doi:10.1021/ct700119m.

44. Kruger, R.; Kuhn, W.; Muller, T.; Woitalla, D.; Graeber, M.; Kosel, S.; Przuntek, H.; Epplen, J.T.; Schols, L.; Riess, O. Ala30Pro mutation in the gene encoding alpha-synuclein in Parkinson’s disease. Nature genetics 1998, 18, 106–108, doi:10.1038/ng0298-106.

45. Kotzbauer, P.T.; Giasson, B.I.; Kravitz, A.V.; Golbe, L.I.; Mark, M.H.; Trojanowski, J.Q.; Lee, V.M.Y. Fibrillization of α-synuclein and tau in familial Parkinson’s disease caused by the A53T α-synuclein mutation. Experimental Neurology 2004, 187, 279–288, 10.1016/j.expneurol.2004.01.007.

46. Kabsch, W.; Sander, C. Dictionary of protein secondary structure: Pattern recognition of hydrogen-bonded and geometrical features. Biopolymers 1983, 22, 2577–2637, 10.1002/bip.360221211.

47. Zhang, Z.; Kang, S.S.; Liu, X.; Ahn, E.H.; Zhang, Z.; He, L.; Iuvone, P.M.; Duong, D.M.; Seyfried, N.T.; Benskey, M.J.;, et al. Asparagine endopeptidase cleaves α-synuclein and mediates pathologic activities in Parkinson’s disease. Nature structural & molecular biology 2017, 24, 632–642, doi:10.1038/nsmb.3433.

48. Braun, A.R.; Lacy, M.M.; Ducas, V.C.; Rhoades, E.; Sachs, J.N. alpha-Synuclein’s Uniquely Long Amphipathic Helix Enhances its Membrane Binding and Remodeling Capacity. The Journal of membrane biology 2017, 250, 183–193, doi:10.1007/s00232-017-9946-1.

49. Vermaas, J.V.; Tajkhorshid, E. Conformational heterogeneity of alpha-synuclein in membrane. Biochimica et biophysica acta 2014, 1838, 3107–3117, doi:10.1016/j.bbamem.2014.08.012.

50. Betarbet, R.; Sherer, T.B.; MacKenzie, G.; Garcia-Osuna, M.; Panov, A.V.; Greenamyre, J.T. Chronic systemic pesticide exposure reproduces features of Parkinson’s disease. Nature Neuroscience 2000, 3, 1301–1306, doi:10.1038/81834.

51. Aksimentiev, A.; Schulten, K. Imaging α-Hemolysin with Molecular Dynamics: Ionic Conductance, Osmotic Permeability, and the Electrostatic Potential Map. Biophysical Journal 2005, 88, 3745–3761, 10.1529/biophysj.104.058727.

52. D.A. Case, R.M.B., D.S. Cerutti, T.E. Cheatham, III, T.A. Darden, R.E. Duke, T.J. Giese, H. Gohlke,A.W. Goetz, N. Homeyer, S. Izadi, P. Janowski, J. Kaus, A. Kovalenko, T.S. Lee, S. LeGrand, P. Li, C.Lin, T. Luchko, R. Luo, B. Madej, D. Mermelstein, K.M. Merz, G. Monard, H. Nguyen, H.T. Nguyen, I.Omelyan, A. Onufriev, D.R. Roe, A. Roitberg, C. Sagui, C.L. Simmerling, W.M. Botello-Smith, J. Swails,R.C. Walker, J. Wang, R.M. Wolf, X. Wu, L. Xiao and P.A. Kollman (2016) *AMBER* 2016, University of California, San Francisco.

53. Huang, J.; Rauscher, S.; Nawrocki, G.; Ran, T.; Feig, M.; de Groot, B.L.; Grubmüller, H.; MacKerell, A.D. CHARMM36m: an improved force field for folded and intrinsically disordered proteins. Nature Methods 2017, 14, 71–73, doi:10.1038/nmeth.4067.

54. Hopkins, C.W.; Le Grand, S.; Walker, R.C.; Roitberg, A.E. Long-Time-Step Molecular Dynamics through Hydrogen Mass Repartitioning. J. Chem. Theory Comput. 2015, 11, 1864–1874, doi:10.1021/ct5010406.

55. Humphrey, W.; Dalke, A.; Schulten, K. VMD: Visual molecular dynamics. Journal of Molecular Graphics 1996, 14, 33–38, 10.1016/0263-7855(96)00018-5.

56. Nguyen, H.; Roe, D.R.; Swails, J.; Case, D.A. PYTRAJ: Interactive data analysis for molecular dynamics simulations., 2016.

57. Harris, C.R.; Millman, K.J.; van der Walt, S.J.; Gommers, R.; Virtanen, P.; Cournapeau, D.; Wieser, E.; Taylor, J.; Berg, S.; Smith, N.J.;, et al. Array programming with NumPy. Nature 2020, 585, 357–362, doi:10.1038/s41586-020-2649-2.

58. Virtanen, P.; Gommers, R.; Oliphant, T.E.; Haberland, M.; Reddy, T.; Cournapeau, D.; Burovski, E.; Peterson, P.; Weckesser, W.; Bright, J.;, et al. SciPy 1.0: fundamental algorithms for scientific computing in Python. Nature Methods 2020, 17, 261–272, doi:10.1038/s41592-019-0686-2.

59. Hunter, J.D. Matplotlib: A 2D Graphics Environment. Computing in Science & Engineering 2007, 9, 90–95, doi:10.1109/MCSE.2007.55.

60. McGibbon, Robert T.; Beauchamp, Kyle A.; Harrigan, Matthew P.; Klein, C.; Swails, Jason M.; Hernández, Carlos X.; Schwantes, Christian R.; Wang, L.-P.; Lane, Thomas J.; Pande, Vijay S. MDTraj: A Modern Open Library for the Analysis of Molecular Dynamics Trajectories. Biophysical Journal 2015, 109, 1528–1532, doi:10.1016/j.bpj.2015.08.015.

