## Supplemental Information for "Molecular Dynamics Study of α-Synuclein Domain Deletion Mutant Monomers"

The center structures for each of the top ten clusters of the isoNAC variant were aligned with respect to the trace and are shown in Figure S1. The overlaid structures show how the NAC domain mostly adopted compact structures when not in contact with the flanking domains.

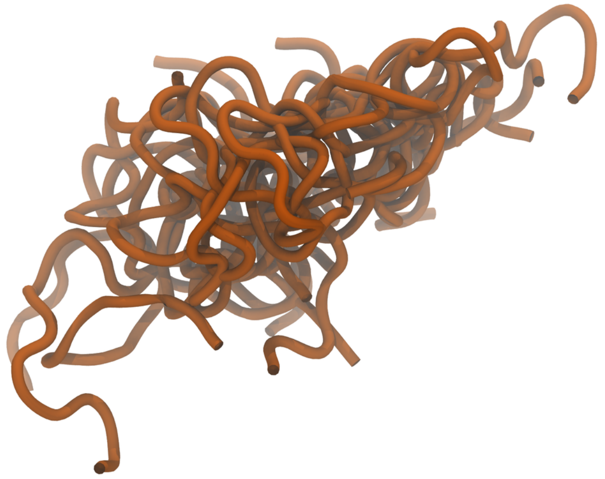

**Figure S1**. The center structures for each of the top ten clusters of the isoNAC variant. The proteins are shown using the New Cartoon representation.

The individual contact maps for the ΔN and ΔC protein variants are shown in Figure S2. The maps show the most probable distance between any pair of residues. Only values up to 12 Å are shown to highlight the contacts.

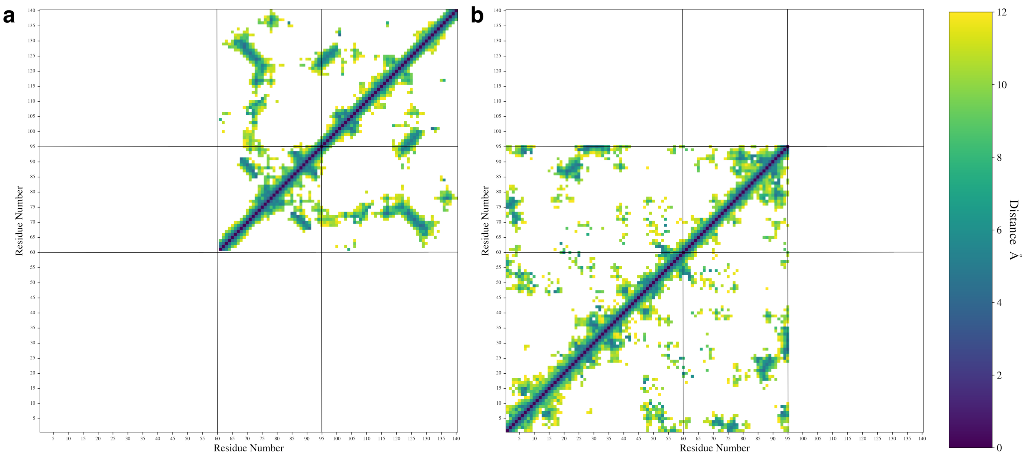


**Figure S2.** The individual contact maps of the most probable distance between amino acid pairs for the ΔN (a), and ΔC (c) variants.

The radius of gyration, $R_g$, of the entire protein is shown for each variant in Figure S3. The $R_g$ generally shows the expected scaling trend, with the smaller isoNAC variant having sampled smaller values and the longer full-length WT protein larger $R_g$ values. The ΔN and ΔC variants, even though their length differs by 15 residues, had similar $R_g$ distributions.

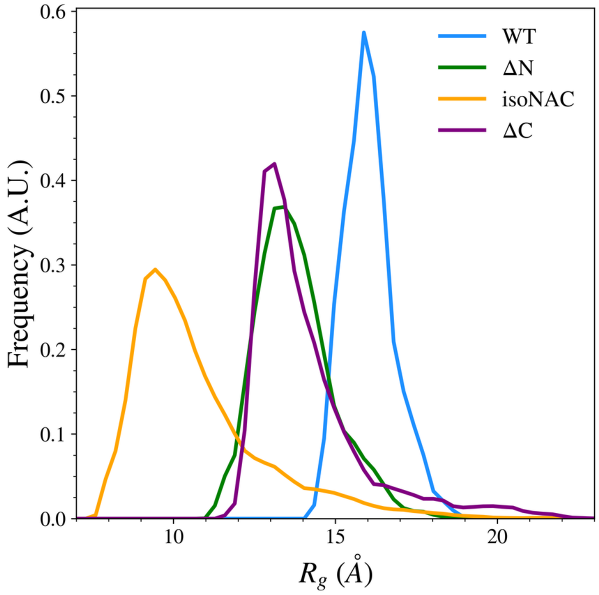


**Figure S3.** The radius of gyration, $R_g$, of the full protein for all simulated variants.
